# Benchmarking of Chromatin Immunoprecipitation (ChIP-Seq) methods in the Pacific oyster *Magallana gigas*

**DOI:** 10.1101/2024.12.28.630603

**Authors:** Amelie Dellong, Manon Fallet, Pierre-Louis Stenger, Cristian Chaparro, Jeremie Vidal-Dupiol, Julie Clément, Céline Cosseau, Christoph Grunau

## Abstract

Chromatin Immunoprecipitation sequencing (ChIP-seq) is the method of choice to generate chromatin landscapes across genomes. The scarcity of literature on ChIPseq and absence of a canonical “gold standard” method in mollusks and especially the pacific oyster *Magallana gigas*, prompted us to compare four ChIP-seq methodologies (Native-ChIP, Crosslink-ChIP, ChIPmentation and Cut & Tag) to find the most suitable method for this species. Our results show that Cut & Tag performs best and ChIPmentation worst. We hypothesize that the reason for this lies in a particularly fragile chromatin structure around genes in the oyster.

## Introduction

Chromatin is a dynamic assembly of DNA and proteins, whose organization and “condensation” levels change to modulate gene function. Post-translational modifications of histones, also known as the “histone code”, are parts of the bearers of epigenetics information (Filion et al., 2010; van Steensel, 2011). We define here epigenetics as the meiotically or mitotically heritable change in gene function that is not based on changes in DNA sequence (Nicoglou and Merlin, 2017). There are different histone modifications which, depending on their nature (such as methylation, acetylation, or phosphorylation) or position on the residues of the N-terminal regions of histones, will influence the activation or repression of gene function (Bannister and Kouzarides, 2011; Barski et al., 2007). The method of choice for mapping these histone marks is “chromatin immunoprecipitation followed by sequencing” or ChIP-seq. This method allows for DNA fragments enrichment linked to a specific protein or, in our case, associated with histone modification. This protocol consists of chromatin fragmentation and the immunoprecipitation of targeted histone marks, the isolation of immunoprecipitated DNA-protein complexes and finally DNA purification, library construction, and sequencing (Barski et al., 2007).

Mollusks are one of the largest and most diverse phyla in the animal kingdom. It brings together numerous species with various major interests: ecological (ecosystem architect such as oyster reefs), health (snail vectors of diseases such as Bilharzia or Fascioliasis), or economic (edible mollusks such as mussels, oysters, etc.). *Magallana gigas*, or the Pacific oyster, is the bivalve species the most used worldwide in aquaculture (Botta et al., 2020; Fallet et al., 2020; Wang et al., 2014). For this species, numerous studies show that epigenetic information is sensitive to environmental variations. DNA methylation, another bearer of epigenetic information, has been studied extensively in *M. gigas* (Fallet et al., 2022; Riviere et al., 2013; Rondon et al., 2017; Sol Dourdin et al., 2023; Venkataraman et al., 2022; Wang et al., 2023, 2021). However, there are very few publications presenting ChIP-seq results in oysters. One publication presents results, in the Pacific oyster, of ChIP-seq concerning HSF1 protein involved in thermal stress response (Liu et al., 2020). Another study on ChIP-seq in *M. gigas* focused on post-translational histone modification. Unfortunately, were unable to reproduce the technical approach and the raw data were inaccessible (Zhan et al., 2015). Therefore, we decided to develop an efficient ChIP-seq protocol in *M. gigas*. As there are many ChIP-seq derivatives, we chose to compare four different ChIP-seq techniques to establish which one is best suited for our animal model.

These four methods are Native ChIP-seq (N-ChIP), Crosslink ChIP-seq (X-ChIP), ChIPmentation and Cleavage Under Targets and Tagmentation (Cut & Tag) (Fig. 1).

**Fig. 1.**
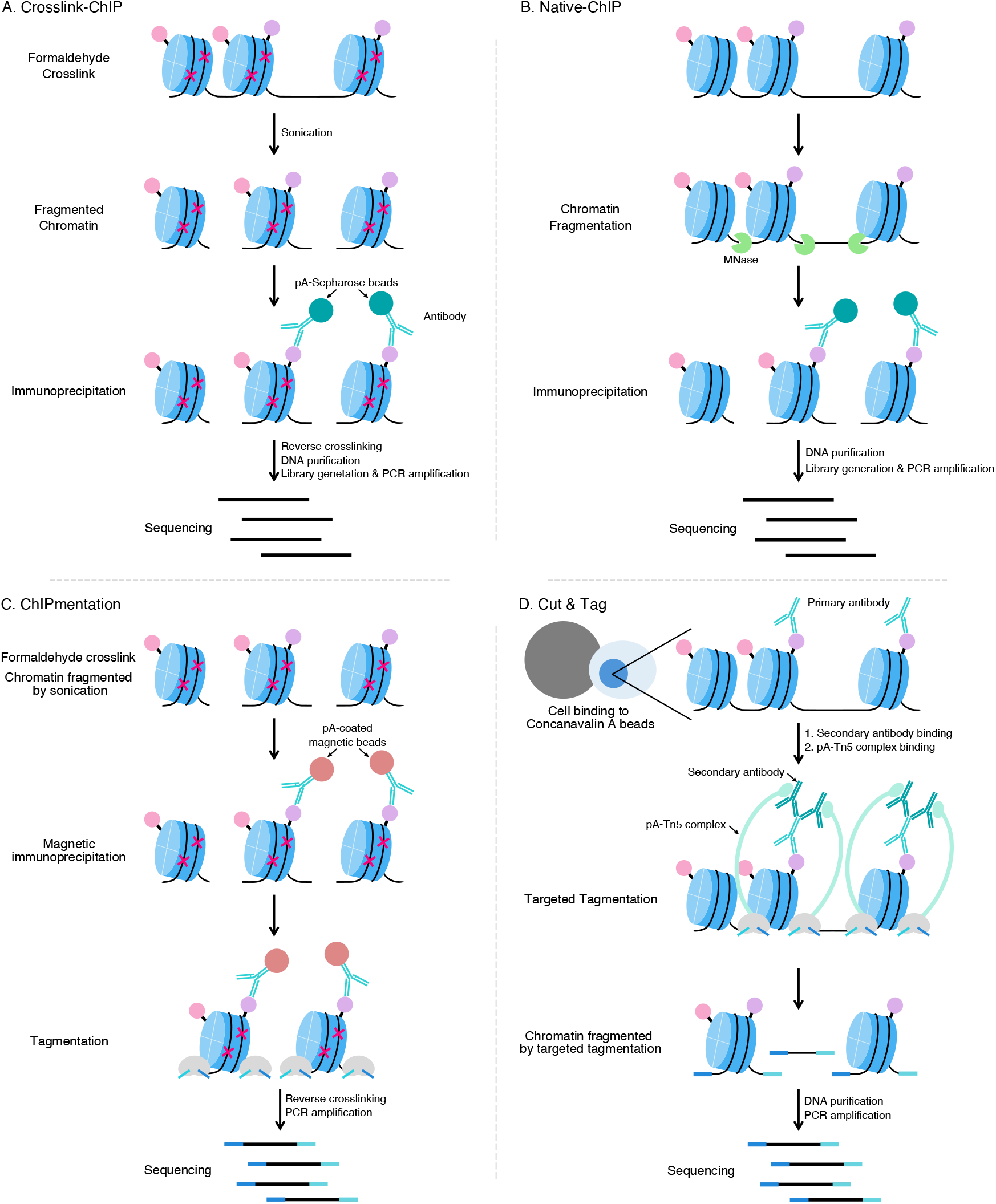
Summary of the four ChIP-seq methods protocols. (A) Crosslink-ChIP, (B) Native-ChIP, (C) ChIPmentation, (D) Cut & Tag. Nucleosomes are represented in blue, crosslinking is represented by fuchsia-colored cross, two different histone modifications are linked to nucleosomes and represented in pink and purple.

N-ChIP required the use of Micrococcal nuclease for chromatin fragmentation. In X-ChIP before the fragmentation step, a crosslinking by adding formaldehyde is required. This step allows the formation of covalent link between DNA and proteins to create fixed complexes. Mechanical chromatin fragmentation is performed by sonication, and before DNA purification it is required to add a crosslink reverse step. For these two methods, after fragmentation, immunoprecipitation with an antibody against a specific histone mark is performed. DNA is purified from the histone-DNA complexes isolated by immunoprecipitation (figure 1A et B) (Park, 2009).

ChIPmentation was published in 2015, it is similar to the X-ChIP protocol but a tagmentation (fragmentation by Tn5) reaction is added after the immunoprecipitation step (Schmidl et al., 2015). Tagmentation used a transposome complex composed of a hyperactive Tn5 transposase and sequencing-compatible adaptors. Tn5 transposase is an enzyme that catalyzes the movement of transposons to other genome regions (Buenrostro et al., 2013). In the ChIPmentation tagmentation reaction, Tn5 fragmented chromatin and insert adapters in bound-beads chromatin simultaneously. In ChIPmentation there are two chromatin fragmentation steps, sonication and tagmentation. This protocol does not necessitate DNA purification on columns. We can also notice that with ChIPmentation protocol, complexes isolation is made by using protein A (pA) coated magnetic beads (Figure 1C) (Schmidl et al., 2015; ChIPmentation Kit for Histones, Diagenode Cat# C01011009)).

Cut & Tag was published in 2019, in this protocol neither MNase digestion, nor crosslinking and sonication are used. Instead, cells or nuclei (if nuclei extraction is performed) must be fixed on a support which are Concanavalin A magnetic beads. All the subsequent reactions are performed in a unique test tube. First, cells are permeabilized and incubated with a primary antibody directed against the targeted histone mark. Then, cells are incubated with a secondary antibody which binds to the primary antibody. Next, the pA-Tn5 complex binds the secondary antibody, this complex allows a targeted tagmentation reaction around the antibody binding site. The binding of the pA-Tn5 complex on the secondary antibody allows for a chromatin fragmentation and adaptors insertion at primary antibody target sites. Finally, tagmented DNA is purified and amplified for sequencing (Figure 1D) (Kaya-Okur et al., 2019; iDeal CUT&Tag kit for Histones, Diagenode Cat# C01070020).

To determine the method best suited for *Magallana gigas*, we decided to work on trimethylated lysine 4 of histone 3 (H3K4me3), one of the most studied marks in other species. In most species, H3K4me3 is associated with open chromatin and to gene transcriptionally active, its signal is mostly located in promoters near the TSS site (Ho et al., 2014; Schuettengruber et al., 2007). It was first necessary to test the effectiveness of commercial antibodies against H3K4me3 in the Pacific oyster. Once the antibody was selected, we compared the four techniques, N-ChIP, X-ChIP, ChIPmentation and Cut & Tag by using metagene profiles around the transcript regions, and input to enrichment ratio. In our hands, Cut & Tag performed best in *M. gigas*.

## Results

### Antibody Ab8580 (Abcam) can be used for Immunoprecipitation of H3K4me3 in M. gigas

Commercial antibodies are available for H3K4me3 but none of them have been tested for oyster histones. Antibody specificity is essential for successful ChIP, and we had previously established a testing pipeline (Cosseau et al., 2009). Briefly, antibodies are first verified by Western Blot where they should deliver a single band around 17kDa and then used the antibody for ChIP titration with increasing amounts of antibody and constant chromatin quantity. Here, an asymptotic increase of input recovery is expected, indicating saturation of targets by the antibody. Only if these two criteria are satisfied, we consider the antibody suitable. Three different antibodies which target H3K4me3, the Abcam Ab8580, the Diagenode C15410003 and the Millipore CS200554/17-678 were tested by Western blot. We expect a unique band on the membrane, at 17kD. For the Abcam and the Diagenode antibodies we observed one unique band between 15 and 20kDa (*figure 1A and 1B*), for the Millipore antibody the signal is located near 15kDa (*figure 1B*), satisfying the criteria 1.

Native ChIP was then performed at an increasing concentration of antibody. For specific antibody-histone interactions, a signal saturation is expected and indicates the quantity of antibodies to be used. For the Diagenode and Millipore H3K4me3 antibody, we did not reach the point where the signal is saturated. (*figure 1D and 1E*). For the Abcam antibody, saturation of IP was obtained using 4µl of antibody (*figure 1C*). Consequently, the Abcam antibody was deemed suitable for ChIP-seq experiments with *M. gigas* histones. We used 4μl of the Abcam antibody at 1g/L for ChIPmentation, X-ChIP and N-ChIP and as a primary antibody in the Cut & Tag experiment.

### Metagene profiles are dependent on the ChIP methods

After having firmly established which antibody to use we wondered which ChIP-seq method would deliver the best signal-to-background ratio. Currently, there is no reference method for ChIP-seq in the oyster. We therefore decided to test four methods (X-ChIP, ChIPmentation, N-ChIP and Cut & Tag). We used as decision criteria metagene profiles around protein-coding genes and comparison of input, signal, and background with heatmaps around enrichment peaks.

Metagene profiles transcribe scores in a profile plot over sets of genomic regions allowing to observe the targeting enrichment in a global way in all genomic regions defined. For Cut & Tag, X-ChIP and N-ChIP we obtained the same types of H3K4me3 profiles with a peak at the transcription start site (TSS) and a ditch at the transcription end site (TES) (*figure 3A, 3B and 3D*). Between these two there is a higher plateau for Cut & Tag and N-ChIP (*figure 3B and 3D*) than for X-ChIP. For ChIPmentation H3K4me3 profile is different from those obtained with the three other methods: we observed a rounded shape curve all along the transcribed regions and a ditch at the TSS instead of a peak.

The signal intensities are also different. For the X-ChIP and Cut & Tag the signal is higher. The downward peak reaches approximately 3.25 and 2.75 respectively and the upward peak extends to 5.5 for X-ChIP and near 6 for Cut & Tag. (*figure 2A and 2D*) while for ChIPmentation and N-ChIP the signal is overall lower and only extends between 1.25 and 1.45 and between approximately 2.8 and 3.8 respectively (*figure 3B and 3C*). These results, all similar except for those obtained by ChIPmentation, led us to rule out this method which does not work on the mantle of *M. gigas*.

**Fig. 2.**
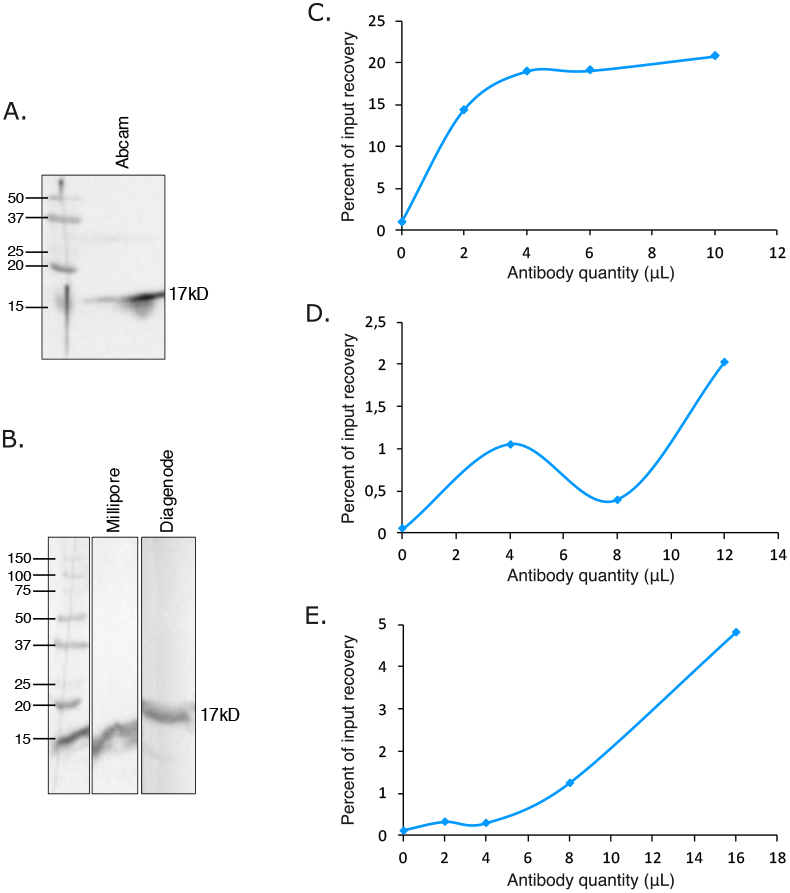
Antibody selection for ChIPseq by western blot and titration. Western blot with the (A) Ab8580 Abcam antibody (B) the Diagenode antibody and the Millipore antibody. qPCR on immunoprecipitated chromatin of *M. gigas* with various antibody volumes for (C) the Abcam, (D) the Diagenode and (E) the Millipore antibody. The amount of target DNA recovered in the immune-precipitated fraction was quantified by calculating the percent input recovery targeting the housekeeping gene encoding the 40S subunit.

**Fig. 3.**
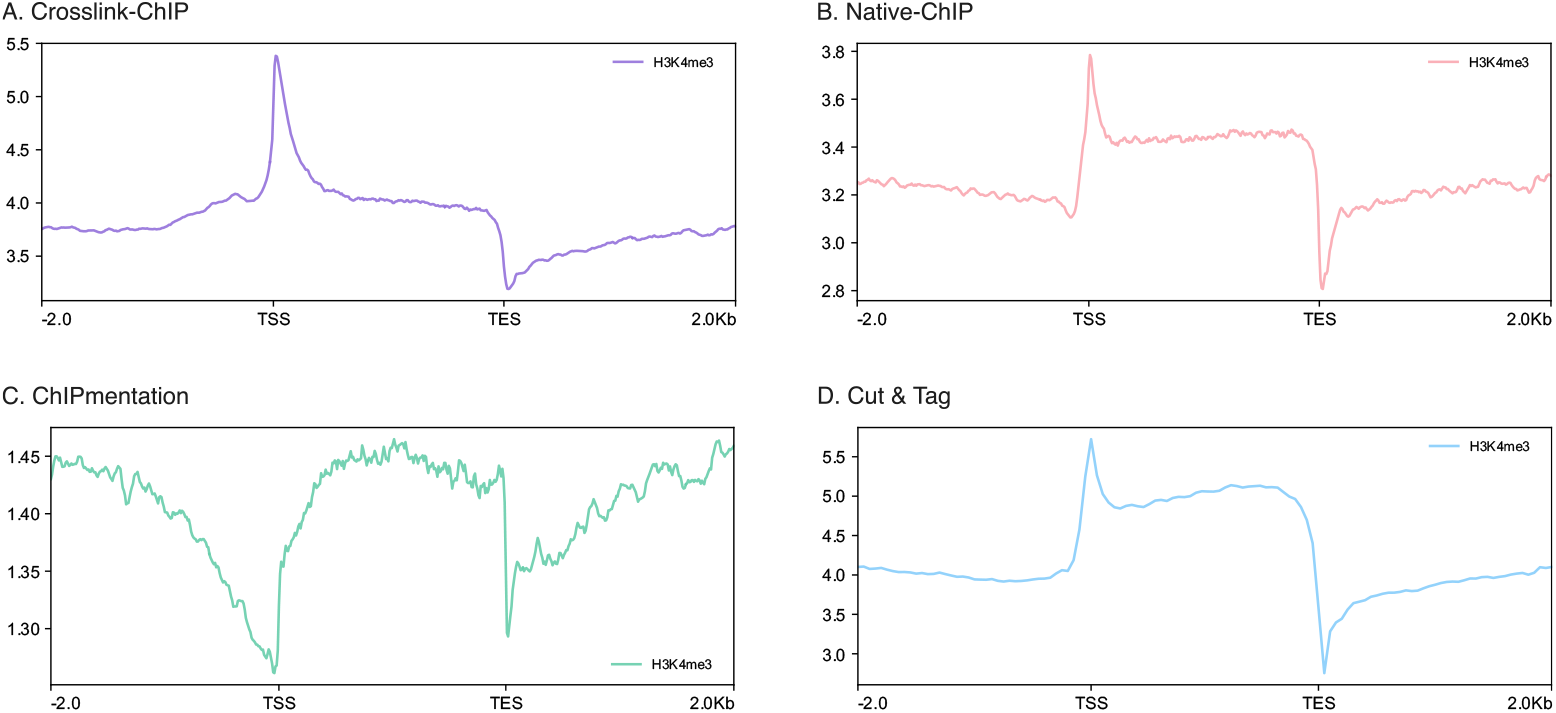
Histone H3K4 tri-methylation in transcribed regions. (A)-(D) H3K4me3 signal intensity (mean of scores attributed by MACS2) between TSS and TES for each experiment with 2kb upstream from TSS and 2kb downstream from TES. Transcript regions are all resized to the same length. (A) for X-ChIP profiles represent a pool of four samples, (B) for ChIPmentation a pool of two samples, for Cut & Tag a pool of four samples and (D) for N-ChIP there is only one sample.

### Cut&Tag has the best signal-to-input ratio

Next, we compared relative enrichment vs background. Enrichment represents a relative measure of H3K4me3 presence compared to the input. While in principle input is the initial amount of chromatin that is used as starting material, each method has different protocols to produce this input which introduces a bias. The background is the amount of chromatin that is precipitated without any antibody. Therefore, it is important to measure input and the background signals.

In X-ChIP, input corresponds to sheared chromatin before immunoprecipitation. In ChIPmentation, we used the same steps performed in X-ChIP but a tagmentation step is added. In this method libraries are therefore fragmented two times. In N-ChIP, chromatin is obtained by digestion with MNase and subsequently incubated with Sepharose-protein-A (pA) beads only. Precipitation delivers the “background” (bound to beads) and the “input” (supernatant available for IP). In contrast to the other three methods, in Cut & Tag, antibody binding precedes chromatin fragmentation, the input i.e. the chromatin available for IP, is not directly accessible. As an alternative, the kit provides an unspecific IgG antibody to generate negative control. In the Cut & Tag experiment, tagmentation is the single fragmentation step and this reaction is directed by the binding of the Tn5 complex to the pA-secondary antibody complex. With an unspecific IgG, no enrichment is expected, and all signal corresponds to the background which must be low in a successful experiment.

To cope with the heterogeneity of the methods in terms of input and background measures we decided to use enrichment profiles and heatmaps centered around peaks for comparison. We considered the best benchmark for specific enrichment to be a high signal to input (for X-ChIP, ChIPmentation and N-ChIP) and signal to IgG as a proxy for input with Cut&Tag. By using Heatmap and Plotprofiles of DeepTools around H3K4me3 centered peaks, it is possible to compare input (or IgG) and ChIP signals. For X-ChIP, ChIPmentation and N-ChIP, the results indicate the presence of an input signal localized at H3K4me3 enrichment sites (*figure 4A, B and C*). The method with the most important input signal on H3K4me3 peaks is the N-ChIP (input signal intensity is stronger than the H3K4me3 signal at H3K4me3 peak locations) (*figure 4B*) followed by ChIPmentation (*figure 4C*) then X-ChIP (*figure 4A*).

**Fig. 4.**
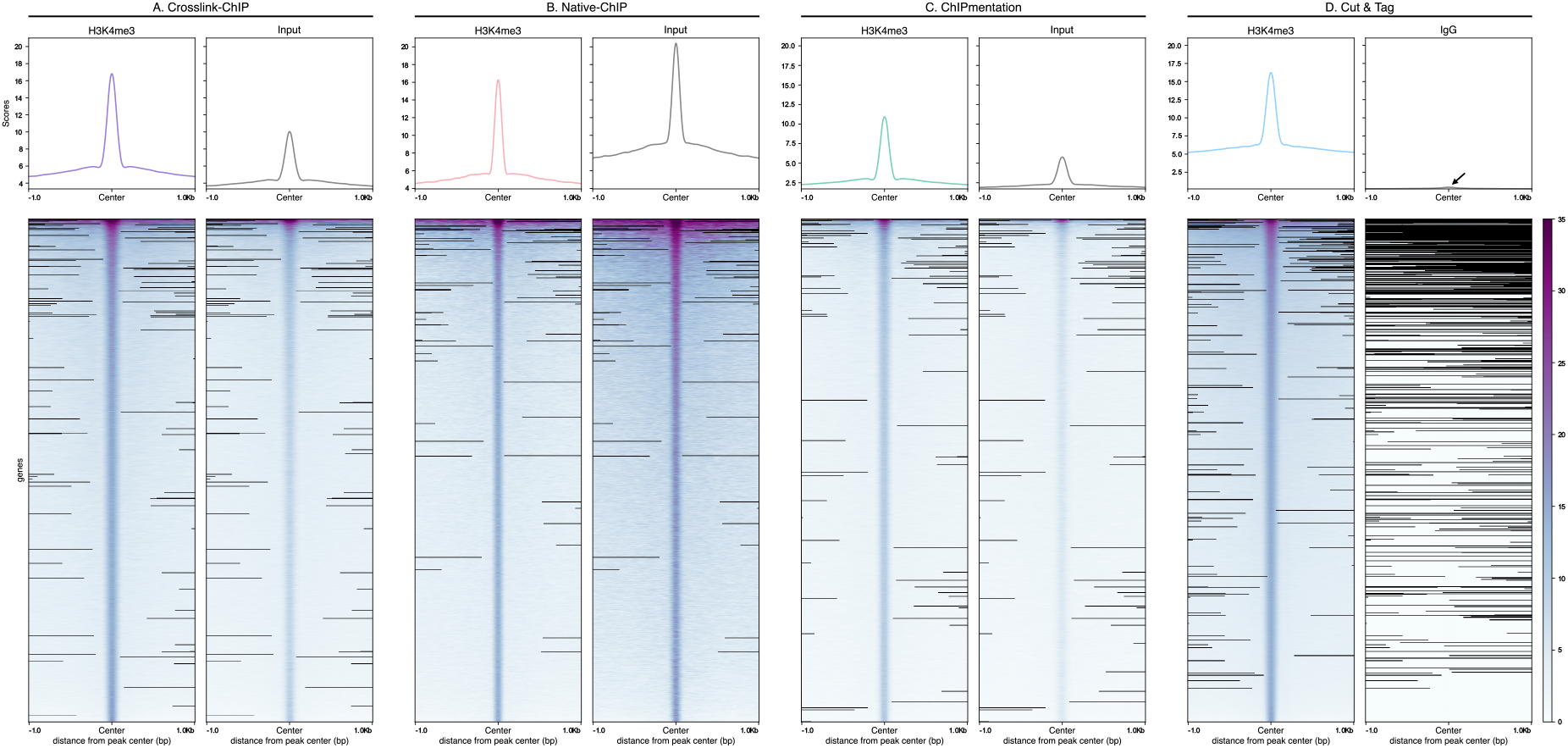
Heatmap centered on peaks for the four ChIP-seq methods end their associate input. (A)-(D) Heatmaps are centered on H3K4me3 peaks with 1kb downstream and upstream from peak center. For each ChIP-seq methods the left panel represent the H3K4me3 signal intensities on the identified H3K4me3 peaks (by MACS2) and the right panel the intensities of input signal on H3K4me3 peaks. H3K4me3 peaks are sorted vertically by descending order and using mean as statistical metric. Black color represents missing data. Metagene profiles are used to plot signal intensity (mean of scores attributed by MACS2) on H3K4me3 peak centers with 1kb downstream and upstream from peak center. (A) for X-ChIP profiles represent a pool of four samples for input and H3K4me3, (B) for N-ChIP there is only one sample for input and H3K4me3 (C) for ChIPmentation a pool of two samples and one sample for input, and (D) for Cut & Tag a pool of four samples and one IgG sample.

In Cut & Tag, we also obtained a peak for the H3K4me3 signal, but the IgG signal is very low with a Heatmap profile totally different from the three inputs and H3K4me3 signals (*figure 4D*). This result corresponds to what is expected for ChIP: it indicates that the obtained signal is specific to the H3K4me3 antibody. In other methods, the input overlaps many H3K4me3 peaks which may indicate the presence of peaks not specific to the antibody and increase the background. For Cut & Tag, the difference between IgG (proxy of input) and H3K4me3 enrichment, is the strongest among the tested methods, reflecting that the signal obtained is specific to H3K4me3 and contains little background noise. We therefore conclude that Cut & Tag is the most suitable method for analyzing of post-translational modifications of histones in the Pacific oyster.

## Discussion

Our results converge towards the conclusion that Cut & Tag is the most appropriate ChIP-seq method for histone post-translational modification analysis in *M. gigas*. Cut & Tag and two other methods (N-ChIP and X-ChIP), gave similar metagene signals spanning transcript regions while the ChIPmentation profile was clearly different. The most parsimonious explanation is that ChIPmentation did not work in the oyster for an unknown reason. Cut & Tag is the method that showed the highest enrichment compared to input. Surprisingly, in N-ChIP, X-ChIP and ChIPmentation already the sequence reads of the input before IP was enriched in the same regions as the H3K4me3 signal. In all of these three methods, chromatin fragmentation precedes IP, and MNase digestion, sonication and tagmentation do not completely randomly fragment the chromatin but target more accessible and therefore open chromatin regions. We hypothesize that *M. gigas* chromatin in the protein coding regions is particularly sensitive to fragmentation methods and therefore already the input signal is strongly present in the H3K4me3 peak locations. The conceptual difference between Cut & Tag and other protocols is that the fragmentation is done after immunoprecipitation. This could explain the improvement of the results with this protocol. An additional advantage of the method is that experiments require hands-on time of only 1.5 days for 12 samples.

While the Cut & Tag method showed superior performance in terms of enrichment compared to the input, one potential limitation of this study is the relatively small number of histone modifications analyzed. Future studies could extend this work to include a broader range of histone modifications and other chromatin-associated proteins to determine whether the observed benefits of Cut & Tag are consistent across different targets.

Obtained results are encouraging regarding the development of Cut & Tag with antibody targeting other histone marks and other oyster tissues. But it is crucial to consider potential biases introduced by the antibody used for immunoprecipitation, as well as the possibility of variation in chromatin accessibility between different tissues within *M. gigas*, or even different species.

The use of Cut & Tag in *M. gigas* not only provides valuable insights into histone post-translational modifications in this species but also holds promise for epigenomic studies in other marine organisms, where traditional ChIP-seq methods may be challenging to implement due to issues with chromatin quality and fragmentation. The ability to efficiently profile histone modifications in non-model species opens up new avenues for comparative epigenomics, particularly in understanding the evolution of chromatin regulation across phylogenetically distant taxa.

## Materials and Methods

### Western blot and antibody titration

Western blot was performed using 100μg of protein extracted from oyster mantle as described previously (Azzi et al., 2009) using the Abcam Ab8580 (lot. GR273043-3), the Diagenode C15410003 (lot. A5051-001P) and the Millipore CS200554/17-678 (lot. NRG1583032). For antibody titration, native ChIP was performed using an increasing amount of H3K4me3 antibodies (for the Abcam 0, 2, 4, 6, 10μl, for the Diagenode 0, 4, 8, 12μl and for the Millipore 0, 2, 4, 8, 16μl). The number of gene-specific sequences associated with each quantity of antibody was quantified by real-time PCR targeting the gene encoding the 40S subunit (Forward primer: aggttttggtggctggattcggt, Reverse primer: agagcgaggagtgatacgttggc, primer Efficiency=2). The percent input recovery of the bound immunoprecipitated fraction for the amplicon was calculated as described previously (Cosseau et al., 2009).

### Native Chromatin Immunoprecipitation (N-ChIP)

300000 pelleted oyster larvae were suspended in 1ml of buffer 1 (0.3M sucrose, 30mM KCl, 7.5mM NaCl, 2.5mM MgCl2, 0.05mM EDTA, 0.1mM PMSF, 0.5mM DTT, 7.5mM Tris–HCl, pH 7.5) containing protease inhibitor cocktail tablets (two tablets for 50ml of buffer 1) (Roche Applied Science, #116974998001) and 5mM sodium butyrate as histone deacetylase inhibitor (Sigma, #B5887). Samples were lysed by adding 1ml buffer 1 with 0.8% NP40 and homogenized in a SZ22 tissue grinder tube (Kontes Glass Company, #885462-0022) using an SC tissue grinder pestle (Kontes Glass Company, #885451-0022) on ice for 7 min. Samples were then processed as described by Cosseau et., al 2009.

Chromatin extracts were quantified, and 30ng of this extract was used for subsequent immunoprecipitation using 4μl of antibody targeting H3K4me3 (Abcam AB8580 lot GR273043-3).

### Cut & Tag

Forty-one – 64 mg of adult oyster mantle tissues were dissected, snap-frozen and stored at -80°C. We then used the Diagenode “Tissue Nuclei Extraction for ATAC-seq” kit (Cat# C01080003, Lot-1) to prepare intact cells which is essential for the Cut & Tag experiment. To lyse the tissue, 800µl of Tissue Lysis Buffer was added directly to a 1.5ml Eppendorf tube and tissue was homogenized using a plastic pestle. Tissue suspension was transferred into a Dounce and further homogenized by 25 strokes with a size A pestle on ice. To remove tissue debris cell suspension was filtered by gravity through a 40μm cell strainer (Sarstedt Cat# 83.3945.040, Lot-12262022) placed on top of a 50ml tube. Flow-through was transferred into new 1.5 ml Eppendorf tubes and centrifuged 5min at 1000g, at room temperature.

We then continued with the Diagenode Cut & Tag Kit (Cat# C01070020, Lot-001, 001A, 001C). Concavalin A beads were prepared according to the suppliers’ instructions, but we then proceeded directly to step 1.14 (Cell Binding). At step 1.16 (resuspension of cells in Complete Wash Buffer 1) we used 1µl for visual inspection of nuclei by adding 5 µl of 1/5000 dilution of DAPI (Invitrogen Cat# D1306, Lot-1891821) in phosphate-buffered saline (PBS) and observation under an epifluorescence microscope at 420 nm and 400x magnification at a NIKON Eclipse TS100 microscope.

To determine the optimal number of amplification cycles for the generation of sequencing libraries we used qPCR. At step 6.2 of the Diagenode protocol, 5µl of the 50µl amplification reaction was withdrawn and we added 0.2µl of SYBR (Invitrogen Cat# S7563, Lot-2415757, diluted 100x). This mixture was used for qPCR on a Mic qPCR Cycler (Bio Molecular Systems) by running the PCR program of step 6.3 (63°C annealing). Optimal cycle numbers were determined by using the number of cycles at which 1/3 of the maximum fluorescence intensity is achieved. Amplification was done with the remaining 45µl following otherwise the Diagenode protocol.

One IgG library was prepared by using the same protocol but with 1ug of IgG antibody provided in kit as primary antibody.

### ChIPmentation

Samples of approximately 20 and 40mg of adult oyster mantle tissues were dissected, snap-frozen and stored at -80°C. For the ChIPmentation protocol we used the Diagenode ChIPmentation Kit (Cat# C01011010 lot-5A, 4B). After the grinding of the samples by plastic pestle in 500μl of HBSS (Cat #14025-050, lot-2470938), 13.5μl of 37% formaldehyde (Sigma-Aldrich Cat# 252549-25mL, Lot-SHBG0805V) were added, and samples were incubated 10min at room temperature for crosslinking reaction. Next, we added 57μl of glycine and samples were incubated for 5 min at room temperature to stop the reaction. After a 5min centrifugation at 500g and at 4°C, cells were resuspended in cold lysis buffer iL1 and a mechanical lysis was performed with a Dounce (A pestle) on ice during 5min, cell suspension is transferred in 1.5ml Eppendorf and centrifuged 5min at 500g and 4°C. Cells were resuspended in cold 1ml iL2 buffer and incubated 10 min at 4°C with rotative agitation. Cells were pellet again with centrifugation 5min at 500g and at 4°C and supernatant was discarded.

Before sonication chromatin samples were transferred into sonication 0.5ml microtubes (Diagenode Cat# C30010020, Lot-20181004). Chromatin shearing was performed by adding 0.5μl of 200x protease inhibitor cocktail and 100μl of shearing buffer iS1 per sample and using the Bioruptor Pico (Diagenode Cat# B01060001) with 5 cycles (one cycle consists of 30 seconds “on”, 30 seconds “off”). These conditions had been previously optimized to obtain fragments of 500 – 1500 bp.

From step 2.9 we followed the Diagenode kit protocol including the additional protocol to perform a sonication test. Magnetic immunoprecipitation and tagmentation were realized on an automated pipetting system (Diagenode SX-8G IP-Star Compact) The optimal number of amplification cycles determination step is also included in the Diagenode protocol.

One Input library was prepared as described in [https://wellcomeopenresearch.org/articles/7-133]. Steps until chromatin shearing are identical to the description above with one sample of approximately 30mg of mantle dissected and snap frozen. We used 1μl of sheared chromatin with 1μl non-diluted Tn5 (Illumina Cat# 15027865, lot-20444992), 10μl 2x tagmentation buffer and 8μl nuclease-free water of molecular biology grade (Sigma-Aldrich Cat# W4502-1L, lot-RNBC0672). Tagmentation buffer was prepared as follows: 20mM Tris(hydroxymethyl)aminoethane, 10mM MgCl2, 20% (vol/vol) dimethylformamide (Wang et al., 2013). The sample was incubated for 5 min at 55°C for tagmentation. Then, 25μl of 2x NEB Next High Fidelity (Cat# M0544S, Lot-10180442) and 5μl MgCl2 (from Diagenode ChIPmentation kit) were added. End-repair and de-crosslink were performed by incubation for 5 minutes at 72°C followed by 10 minutes at 95°C. Finally, for PCR amplification 2.5µl of forward universal ATAC-seq primer (25µM) and reverse ATAC-seq primers (Buenrostro et al., 2013) (25µM) were added.

The qPCR mix for determination of optimal amplification cycle number was prepared as described in the reference protocol (Diagenode Cat# C01011010) except for SYBR, we used a 10x dilution of SYBR (Invitrogen Cat# S7563, Lot-2415757). We used Mic qPCR Cycler for qPCR by running the PCR program of step 5.4 of the Diagenode ChIPmentation protocol (63°C annealing).

### Crosslink Chromatin Immunoprecipitation (X-ChIP)

Fresh gills piece of 100mg was dissected from adult oysters and subdivided in 4 pieces. X-ChIP was performed according to the Diagenode “Auto iDeal ChIP-seq kit for histones” (Cat# C01010057) manual. This protocol required the use of the Diagenode SX-8G IP-Star Compact. Sonication was performed with Diagenode Bioruptor Sonication System UCD-200TM-EX Lab with 3 cycles (one cycle 5min with the 30s “on”, 30s “off” 5 times). Again, these conditions had been established to obtain fragments of 500 – 1500 bp. For immunoprecipitation 4μg of antibody Ab8580 (Abcam, lot-GR273043-3) was used.

### Determination of PCR amplification rounds

To determine the optimal cycle number for library amplification we used the fluorescence curve obtained by qPCR. When the fluorescence curve reaches the plateau, we used a third of the associated value and by projection on the curve we determined the optimal number of cycles for library amplification.

### Sequencing and Bioinformatics analysis

After PCR amplification libraries were purified with AMPure. Quality control and quantification were performed with Agilent Bioanalyzer 2100 Instrument and Agilent High Sensitivity DNA Kit (Cat# 5057-4626). Libraries were sequenced on an Illumina NextSeq550. All data treatments were carried out under a local galaxy instance (http://bioinfo.univ-perp.fr).

For Cut & Tag and ChIPmentation, no adaptor trimming was necessary, and reads were directly aligned to the *M. gigas* genome Roslin v1 (Peñaloza et al., 2021) using Bowtie2 (Galaxy Version 2.3.4.3 + galaxy0) with default parameters. For peak calling we used MACS2 (Galaxy Version 2.1.1.20160309.6) with default parameters and with an effective genome size of 600 Mb. For N-ChIP and X-ChIP, we used the same bioinformatics pipeline but before alignment, we added reads quality filtering and adapter trimming performed with TrimGalore (Galaxy Version 0.6.3) with default parameters except for “phred quality score threshold” for which we used 26.

After using MACS2 for peak calling, the bedgraph files obtained were converted into bigwig files with the converter “Convert BedGraph to BigWig” (Galaxy Version 1.0.1). The data preparation for plotting profile and heatmap of the regions of interest was performed with Deeptools computeMatrix (Galaxy Version 3.5.1.0.0).

For plotting of metagene profiles, computeMatrix was used with default parameters, in scale-regions mode and with 2000 as “Distance in bases to which all regions are going to be fit”. For the regions to plot we filtered the genome annotation file to conserve only mRNA regions. Next, to generate the metagene profile we used DeepTools plotProfiles (Galaxy Version 3.5.1.0.0) with default parameters.

For heatmaps, computeMatrix was used with default parameters, in reference point mode and the center of the region as reference point for plotting. For the region to plot file we used the H3K4me3 narrow peaks file obtained in the output of MACS2 for each ChIP-seq method. To generate the heatmaps we used DeepTools plotHeatmaps (Galaxy Version 3.5.1.0.1) with default parameters.

## Author Contributions and Notes

C.C. and C.G. designed research, A.L., M.F. and P-L.S. performed research, C.C. adapted software, all authors analyzed data and wrote the paper.

The authors declare no conflict of interest.

## Acknowledgments

A.D holds a PhD grant of the Région Occitanie and of the UPVD. M.F held a PhD grant of the ED305 of the university of Perpignan.

This study is set within the framework of the « Laboratoire d’Excellence (LabEx) » TULIP (ANR-10-LABX-41). We thank the Bio-Environment platform (University of Perpignan Via Domitia) and Jean-François Allienne for support in library preparation and sequencing. With the support of LabEx CeMEB, an ANR « Investissements d’avenir » program (ANR-10-LABX-04-01) and the Environmental Epigenomics Core Service at IHPE.

## Notes

### Competing Interest Statement

The authors have declared no competing interest.

